# DNA Methylation Signatures of a Large Cohort Monozygotic Twins Clinically Discordant for Multiple Sclerosis

**DOI:** 10.1101/381822

**Authors:** Nicole Y. Souren, Lisa A. Gerdes, Pavlo Lutsik, Gilles Gasparoni, Eduardo Beltran, Abdulrahman Salhab, Tania Kümpfel, Dieter Weichenhan, Christoph Plass, Reinhard Hohlfeld, Jörn Walter

## Abstract

Multiple sclerosis (MS) is an inflammatory demyelinating disease of the central nervous system with a modest concordance rate in monozygotic twins that strongly argues for involvement of epigenetic factors. We observe in 45 MS discordant monozygotic twins highly similar peripheral blood mononuclear cell-based methylomes. However, a few MS-associated differentially methylated positions (DMP) were identified and validated, including a region in the *TMEM232* promoter and *ZBTB16* enhancer. In CD4+ T cells we observed an MS-associated differentially methylated region in *FIRRE.* In addition, many regions showed large methylation differences in individual pairs, but were not clearly associated with MS. Furthermore, epigenetic biomarkers for current interferon-beta treatment were identified, and extensive validation revealed the *ZBTB16* DMP as a signature of prior glucocorticoid treatment. Altogether, our study represents an important reference for epigenomic MS studies. It identifies new candidate epigenetic markers, highlights treatment effects and genetic background as major confounders, and argues against some previously reported MS-associated epigenetic candidates.

---

Multiple sclerosis (MS) is one of the leading causes of neurological disability in young adults^1^, and is considered to be an autoimmune disease, characterized by chronic inflammatory demyelination of neurons in the central nervous system^2^. Although nuclear genetic factors contribute to the development of MS^3^, the maximum reported concordance rate for MS in monozygotic (MZ) twins is only 25%^4,5^, which indicates that for the development of clinical symptoms interaction with other risk factors remains compulsory. Despite of various studies suggesting that mitochondrial DNA variants are plausible susceptibility factors for MS, we recently showed that mitochondrial DNA variation (e.g. skewed heteroplasmy) does not play a major role in the discordant clinical manifestation of MS in MZ twins^6^.

Another source of molecular variation that can cause discordant phenotypes within MZ twins are DNA methylation differences^7-10^. DNA methylation is an epigenetic modification that in mammals almost exclusively occurs at cytosines within CpG dinucleotides. Since changes in CpG methylation can cause transcriptional alterations, DNA methylation is crucial for normal development and aberrant DNA methylation has been observed in various human diseases^11,12^. Discordant DNA methylation profiles within MZ twins have quite frequently been reported at imprinted regions^7,8^, which are characterized by parent-of-origin-specific methylation patterns resulting in mono-allelic expression. Since a number of studies reported a maternal parent-of-origin effect in MS susceptibility^13,14^, and several imprinted genes have been linked to immune system development and functioning (reviewed by Ruhrmann et al.^15^), genomic imprinting errors might be involved in the pathogenesis of MS^15^. In addition, environmental risk factors such as smoking, history of symptomatic Epstein-Barr virus infection and vitamin D deficiency have been associated with an increased MS risk^16-18^. Although the molecular mechanisms underlying these associations remain unknown, it is possible that these environmental factors operate by inducing DNA methylation changes.

Thus far, several epigenome-wide association studies (EWAS) for MS have been carried out^19-25^, and a number of differentially methylated CpG positions (DMPs) have been reported, including DMPs in the *HLA-DRB1* locus. However, even though these studies used the same array platform (i.e. Infinium HumanMethylation450 (450K) array), the reported results are inconsistent. Since these studies used genetically unmatched cases and controls, they are potentially hampered by DNA sequence variation. As genetic factors predispose to MS, these studies cannot dissect whether MS is due to genetic or epigenetic susceptibility. In addition, SNP-containing probes give rise to biased DNA methylation measurements^26^, and DNA methylation changes are also often the result of *cis-* or *trans-acting* genetic variants (methylation quantitative trait loci or mQTLs)^27^. Fortunately, an MZ twin-based design controls for these genetic differences as well as for other factors (potentially) affecting the methylome, including gender, age and a broad range of environmental factors. Thus far, one EWAS in MS discordant MZ twins has been reported, in which no DNA methylation differences were identified^28^. Since this study included only three twin pairs, further studies in larger cohorts are required. Hence, here we describe an EWAS comprising a unique cohort of 45 MZ twins clinically discordant for MS, in which we aim to identify DNA methylation changes associated with the clinical manifestation of MS and to study the effect of MS treatments on the DNA methylome (**Fig. 1**).

**Figure 1.**
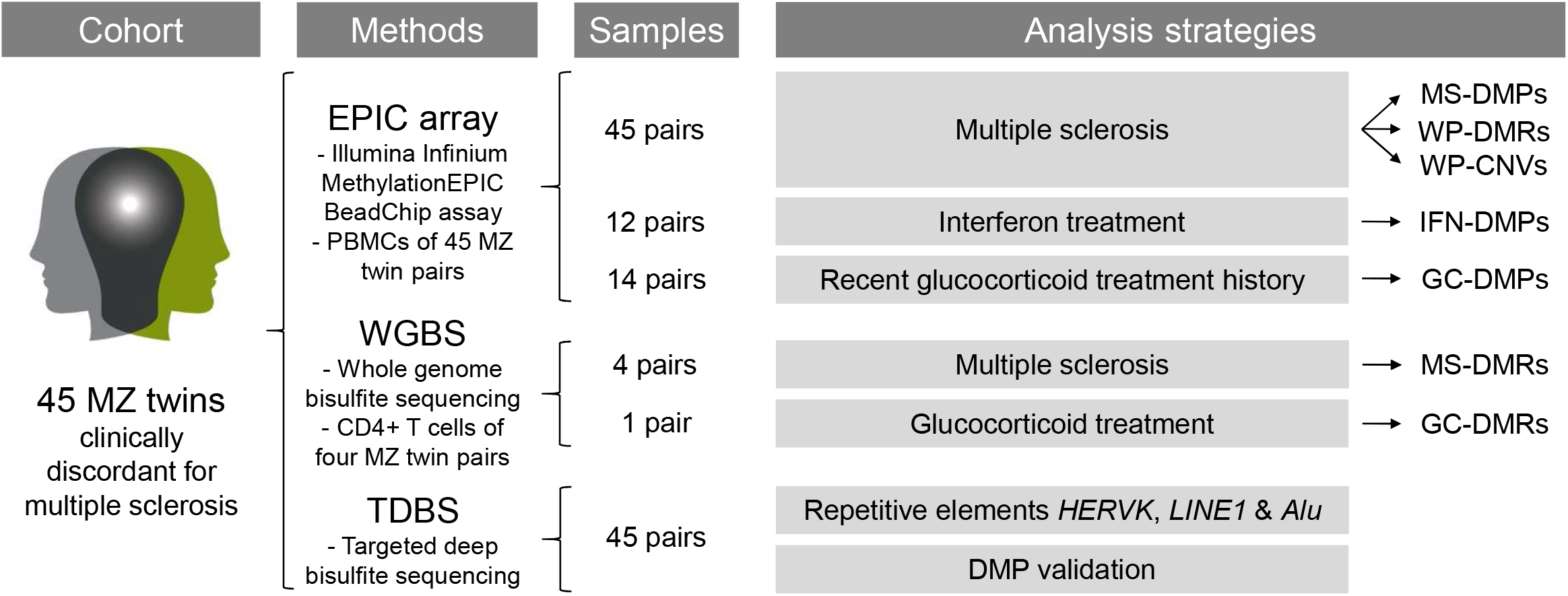
Schematic overview of the study design and analysis strategies. DMPs, differentially methylated CpG positions; DMRs, differentially methylated regions; GC-DMPs, glucocorticoid treatment-associated DMPs; HERVK, human endogenous retrovirus type K; IFN-DMPs, interferon treatment-associated DMPs; LINE1, long interspersed nuclear element-1; MS-DMPs, multiple sclerosis-associated DMPs, MZ, monozygotic; PBMCs, peripheral blood mononuclear cells; WP-CNVs, within-pair copy number variations; WP-DMRs, within-pair differentially methylated regions.

## Results

### Peripheral blood mononuclear cell-based methylomes

For this study peripheral blood mononuclear cells (PBMCs) of 46 MZ twins clinically discordant for MS were accessible and genome-wide DNA methylation profiles were established using Illumina Infinium MethylationEPIC BeadChips (EPIC arrays). After quality control and filtering, methylation data of 849,832 sites were available for 45 MZ twin pairs. As expected, these twin pairs showed a very high mean within-pair array-wide correlation coefficient of 0.995, indicating high quality data. General demographic and clinical characteristics of these 45 MS discordant MZ twins at study entry are shown in **Table 1**.

**Table 1.**
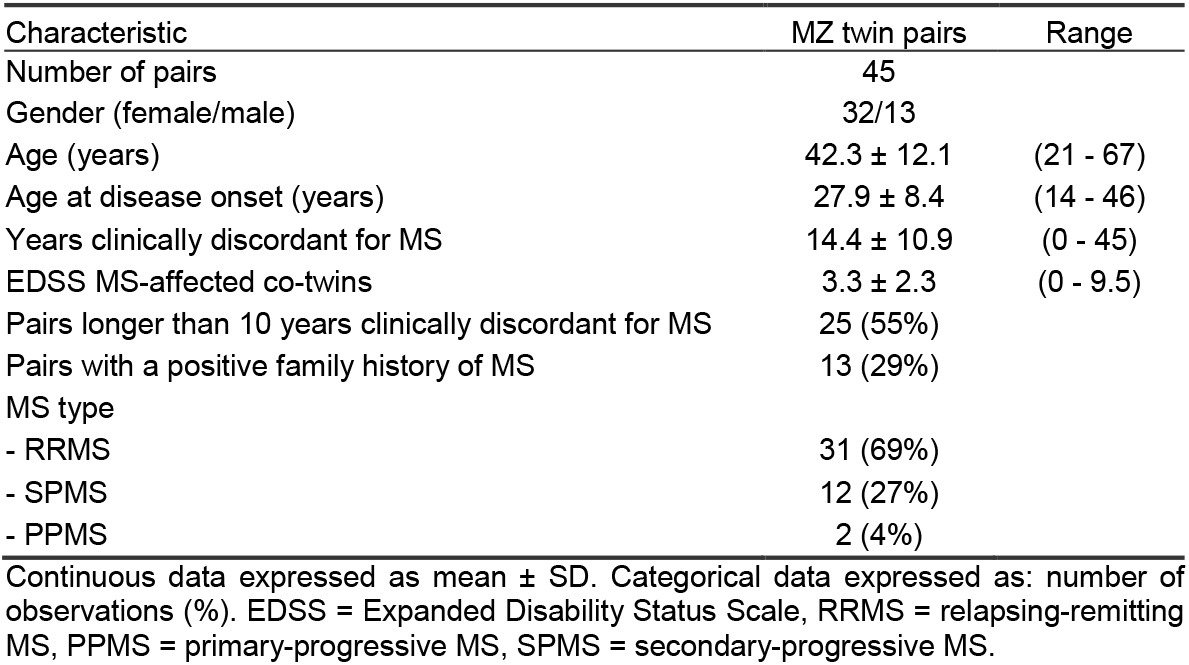
General demographic and clinical characteristics of the MZ twins clinically discordant for MS at study entry.

| Characteristic | MZ twin pairs | Range |
| --- | --- | --- |
| Number of pairs | 45 |  |
| Gender (female/male) | 32/13 |  |
| Age (years) | 42.3 ± 12.1 | (21 - 67) |
| Age at disease onset (years) | 27.9 ± 8.4 | (14 - 46) |
| Years clinically discordant for MS | 14.4 ± 10.9 | (0 - 45) |
| EDSS MS-affected co-twins | 3.3 ± 2.3 | (0 - 9.5) |
| Pairs longer than 10 years clinically discordant for MS | 25 (55%) |  |
| Pairs with a positive family history of MS | 13 (29%) |  |
| MS type |  |  |
| - RRMS | 31 (69%) |  |
| - SPMS | 12 (27%) |  |
| - PPMS | 2 (4%) |  |

Continuous data expressed as mean ± SD. Categorical data expressed as: number of observations (%). EDSS = Expanded Disability Status Scale, RRMS = relapsing-remitting MS, PPMS = primary-progressive MS, SPMS = secondary-progressive MS.

### Identification and confirmation of MS-associated DMPs in *TMEM232* and *ZBTB16*

To identify DMPs associated with the clinical manifestation of MS (MS-DMPs), first a pair-wise analysis was carried out on the EPIC array data of the 45 pairs without adjusting for cell type composition (**Supplementary Fig. 1a** and **1b**). The Q-Q plot in **Supplementary Fig. 1b** shows that the obtained P-values clearly deviate from the null expectation. This inflation of P-values was eliminated after adjusting for cell type composition (**Fig. 2a** and **2b**), indicating that many differences are due to variation in cellular composition.

**Figure 2.**
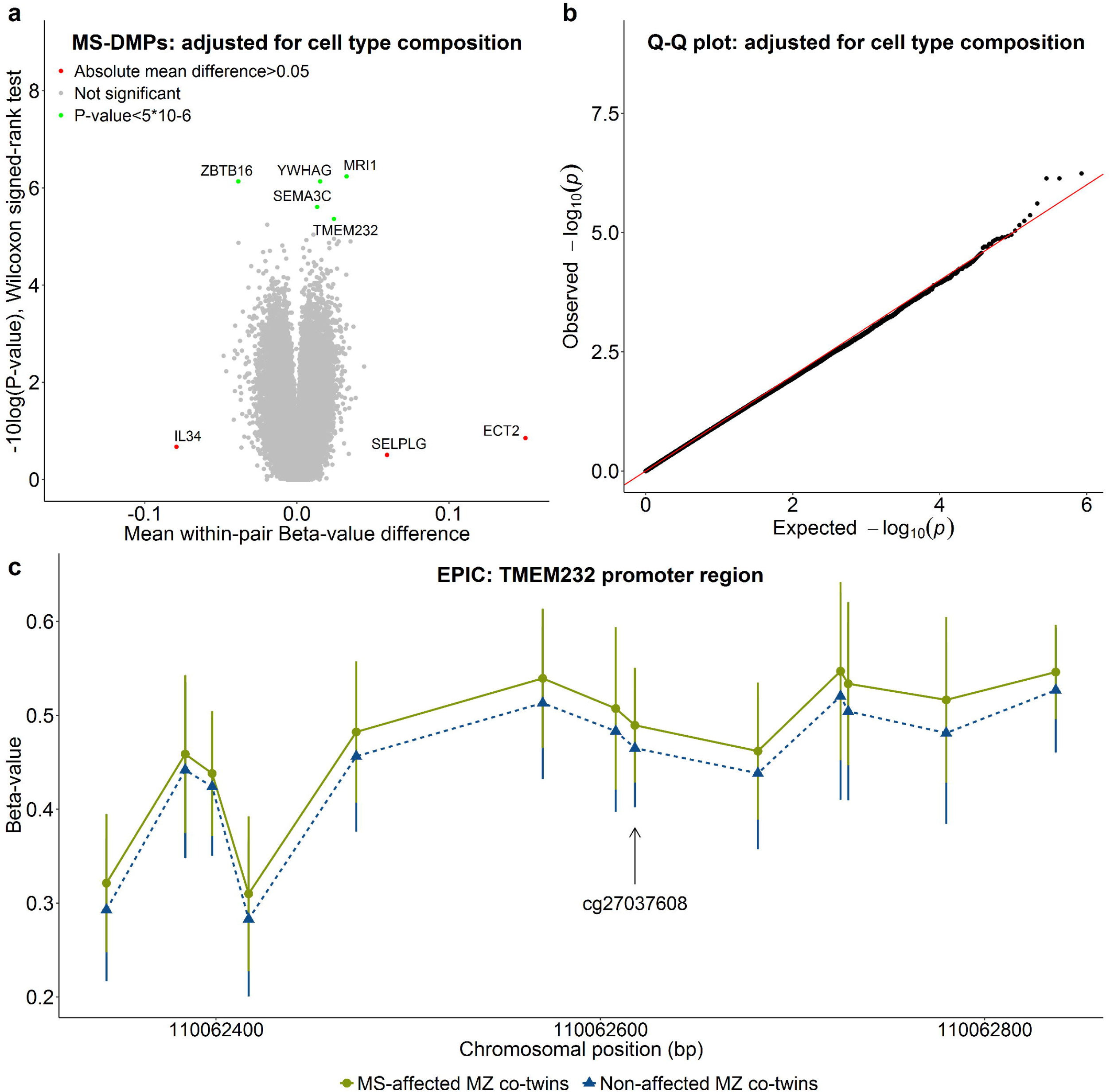
DNA methylation changes associated with the clinical manifestation of MS. Results of the differential DNA methylation analysis including the EPIC array data of the 45 MZ twin pairs clinically discordant for MS. (**a**) Volcano plot of the P-values resulting from the non-parametric Wilcoxon signed-rank test against the mean within-pair β-value difference for each CpG. Data were adjusted for cell-type composition. (**b**) Q-Q plot of the P-values resulting from the non-parametric Wilcoxon signed-rank shown in Fig. 3a. Data were adjusted for cell-type composition. Within-pair β-value difference (Δβ-value) = clinically MS-affected MZ co-twin – nonaffected MZ co-twin. (c) Overview of the *TMEM232* promoter region, including the mean (adjusted) β-values of the MS-affected and clinically non-affected MZ co-twins of the significant MS-associated differentially methylated CpG position (MS-DMP) cg27037608 and 12 neighbouring CpGs present on the EPIC array.

The volcano plots in **Fig. 2a** and **Supplementary Fig. 1a** show that the mean within-pair β-value differences (Δβ-values) are small. The largest differences are observed for *ECT2* (cg12393503), *SELPG* (cg02520593) and *IL34* (cg01447350), with mean Δβ-values of 0.15, 0.06 and −0.09, respectively, but with non-significant P-values. In several twins these CpGs show very large Δβ-values (up to 0.8) (**Supplementary Fig. 2**). However, these large within-pair methylation differences were not confirmed by validation using targeted deep bisulfite sequencing (TDBS) (**Supplementary Fig. 3**). This indicates that some EPIC probes are prone to technical artefacts, as reported by others^29^, and that validation using independent assays is required.

The unadjusted analysis revealed 39 MS-DMPs with a suggestive P-value<5*10^-6^ and after correcting for multiple testing six MS-DMPs remained genome-wide significant (false discovery rate (FDR)<0.05) (**Supplementary Fig. 1a**). After adjusting for cell-type composition, no MS-DMP had an FDR<0.05, but five MS-DMPs had a suggestive P-value<5*10^-6^ (**Fig. 2a** and **Table 2**). One of these MS-DMPs is located in the promoter of the *TMEM232* gene (cg27037608, mean Δß-value=0.024), encoding for a transmembrane protein of unknown function. However, genetic variants in this gene have been associated with atopic dermatitis and allergic rhinitis in genome-wide association studies (GWAS)^30-32^. For this MS-DMP EPIC array data of 12 neighbouring CpGs were also available, which all showed a similar effect and eight CpGs had a P<0.01 (**Fig. 2c**). A second solitary MS-DMP was observed in the gene body of *SEMA3C* (cg00232450, mean Δß-value=0.013), which has been suggested to promote migration of dendritic cells during innate and adaptive immune responses, and is considered to be involved in axonal guidance and growth^33,34^. A third MS-DMP is located in the *YWHAG* gene (cg01708711, mean Δß-value=0.015), which has been associated with MS severity in a GWAS^35^. This MS-DMP has five neighbouring CpGs on the array, but all with non-significant P-values (>0.01), questioning the significance of this MS-DMP. A fourth MS-DMP (cg25345365) has the largest methylation difference (mean Δß-value=-0.039) and is located in an enhancer in the *ZBTB16* gene, which has been reported to be essential for natural killer T (NKT) cell development^36^. Finally, the fifth MS-DMP (cg25755428, mean Δß-value=0.033) is located in the *MRI1* gene, mutations in which have been associated with vanishing white matter disease^37^. The β-value distribution of this CpG and the neighbouring ones in the twins, suggest that this MS-DMP concerns an mQTL (**Table 2**).

**Table 2.**
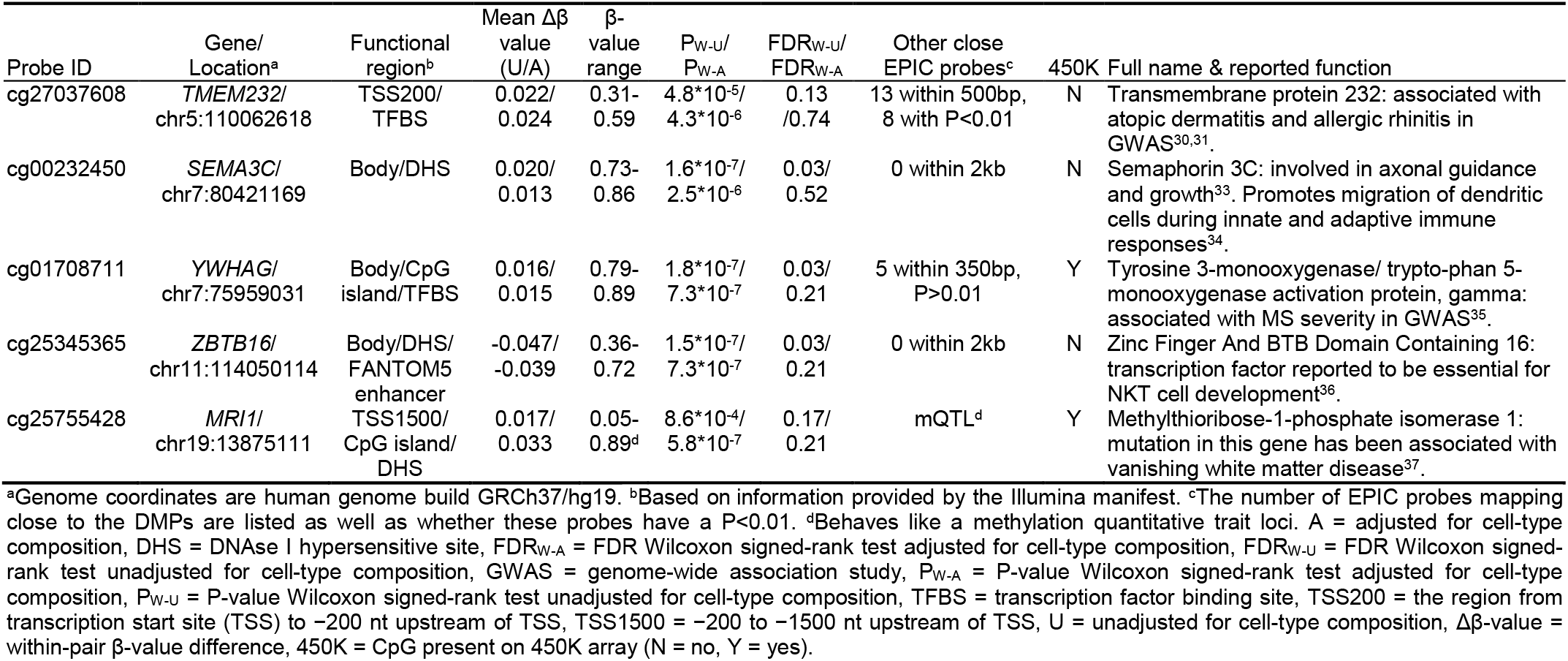
DMPs associated with the clinical manifestation of MS (MS-DMPs). Listed are the five MS-DMPs with a suggestive P-value<5*10^-6^ in the pair-wise analysis carried out using the EPIC array data of the 45 MZ twin pairs adjusted for cell-type composition.

Next, a functional annotation analysis carried out with all MS-DMPs having a P<0.001 in the analysis adjusted for cell-type composition (698 in total) revealed *TMEM232* as a prime candidate marker for MS. All other annotation categories were not significant.

Based on their significance, effect size and/or whether neighbouring probes were also differentially methylated, the *TMEM232* (cg27037608) and *ZBTB16* (cg25345365) MS-DMPs were selected for validation using TDBS. The TDBS data correlated highly with the array data (r**r**_TMEM232_=0.84 and **r**_ZBTB16_=0.89, **Supplementary Fig. 4a** and **5a**). In addition, both the individual MS-DMPs as well as the surrounding CpGs in the *TMEM232* and *ZBTB16* loci were significantly differentially methylated between the clinically non-affected and MS-affected co-twins (**Supplementary Fig. 4b** and **5b**). This confirms that these MS-DMPs represent true effects in our twin cohort.

### Whole genome bisulfite sequencing identifies MS-DMR in *FIRRE*

To identify additional MS-associated differentially methylated regions (MS-DMRs), we performed a pilot study using whole genome bisulfite sequencing (WBGS) in a subset of four MS discordant female twin pairs. Since raw PBMC-based methylomes are hampered by cell type composition differences and need adjustment, we decided to perform WGBS on CD4+ memory T cells, which have been implicated in the pathogenesis of MS^38^. First global DNA methylation changes in the WGBS data were evaluated by identifying and comparing partially methylated domains (PMD), fully methylated regions (FMRs), low methylated regions (LMRs) and unmethylated regions (UMRs). However, no global changes were observed between the non-affected and MS-affected MZ co-twins (P>0.05) (**Supplementary Fig. 6** and **7**).

Next, a DMR analysis was carried out and MS-DMRs were defined as ≥3 CpGs (max. distance 500bp), each having P<0.05 and absolute mean methylation difference >0.2. This analysis revealed a prominent MS-DMR located in an intronic CTCF/YY1 bound regulatory region in the *FIRRE* gene, that is located on the X-chromosome (chrX:130863481-130863509) and encodes a circular long non-coding RNA (**Supplementary Fig. 8**)^39^. The CpGs within this MS-DMR are not covered by the EPIC array, but a probe (cg08117231) located close to this DMR was not significant in the PBMC-based EWAS (P>0.05). The *TMEM232* and *ZBTB16* loci did not fulfill our filtering criterion of ≥10 reads coverage across all samples (note that they were validated by TDBS).

### Within-pair differentially methylated regions are common among MZ twins

The EPIC array-based association analysis enables to identify relatively small DNA methylation differences present in a large number of cases. However, since MS is a heterogeneous disease, DNA methylation changes present in only a few cases might also have biological consequences. We therefore performed a DMR analysis in the individual twin pairs to identify so-called within-pair DMRs (WP-DMR) using the EPIC array data. To detect robust methylation changes, WP-DMRs were defined as ≥3 CpGs within 1 kb having a Δβ-value (adjusted for cell-type composition) >0.20, and the aberrant methylated co-twin having a β-value greater than ±3 standard deviations from the mean. Overall, 45 WP-DMRs were identified in 17 out of the 45 twin pairs, ranging from one to 11 WP-DMRs per pair (**Supplementary Table 1** and **Supplementary Fig. 9-12**). Of the 45 WP-DMRs, 43 were solitary pair-specific and only two were found in two independent pairs. These recurrent WP-DMRs were located in the *ISOC2* and the *HIST1H3E* promoter (**Supplementary Fig. 9**), but for both WP-DMRs the aberrant methylation pattern did not correlate with the MS phenotype (Supplementary Table 1). Of the 43 pair-specific WP-DMRs, 16 showed an aberrant methylation pattern in the non-affected cotwin and 27 in the MS-affected co-twin. Thus far these WP-DMRs have not been associated with MS in other EWAS, neither overlap with MS-associated genes reported in the GWAS Catalog (accessed May 2018)^40^. Two of the pair-specific WP-DMRs were located in reported imprinted regions^41^, including *SVOPL* and *HM13/MCTS2P* (**Supplementary Fig. 10**), but in both cases the non-affected co-twin showed an abnormal methylation pattern. Furthermore, in one pair four WP-DMRs were identified in the protocadherin gamma *(PCDHG)* gene cluster due to hypermethylation in the MS-affected co-twin (**Supplementary Fig. 11**). In another pair a WP-DMR was observed in the promoter of the non-clustered protocadherin 10 *(PCDH10)* gene, also due to hypermethylation in the MS-affected co-twin (**Supplementary Fig. 12**). Protocadherins are highly expressed in the brain and are involved in neuronal development^42^.

### Interferon treatment induces robust DNA methylation changes

Besides identifying methylation changes associated with the clinical manifestation of MS, our MZ twin design also enables to identify MS treatment-related DNA methylation changes. Since in our cohort interferon-beta (IFN) is the most common disease modifying treatment, we performed a pair-wise analysis only including the EPIC array data of the 12 pairs of which the MS-affected co-twins were treated with IFN at the moment of blood collection. We observe that the mean within-pair β-value differences (Δβ-values) for this sub-cohort are larger (**Fig. 3a**), since 257 DMPs with an absolute mean Δß-value>0.05 and P<0.001 were identified. None of the MS-DMPs listed in **Table 1** were among these 257 IFN-associated DMPs (IFN-DMPs). The 257 IFN-DMPs were annotated to 212 genes, of which 124 genes (58%) overlap with IFN-regulated genes recorded in the INTERFEROME database (accessed May 2018), which is based on gene expression data^43^. In addition, a functional annotation analysis revealed a clear enrichment for genes involved in the antiviral defence and interferon homeostasis (**Fig. 3b**). Moreover, seven IFN-DMPs had an absolute mean Δß-value>0.10 and P<0.001, due to strong hypomethylation in the IFN-treated MS-affected co-twins (**Supplementary Table 2** and **Supplementary Fig. 13**). These seven DMPs were located in *RSAD2* (n=3), *MX1* (n=2), *IFI44L* (n=1) and *PLSCR1* (n=1), i.e. genes that have been reported to be up-regulated in blood and PBMCs of MS patients following IFN treatment^43-45^. Adjusting the data for cell-type composition resulted only in a slight attenuation of the IFN-effect (**Supplementary Table 2**), indicating that these seven DMPs are robust markers for monitoring IFN treatment effects in PBMCs.

**Figure 3.**
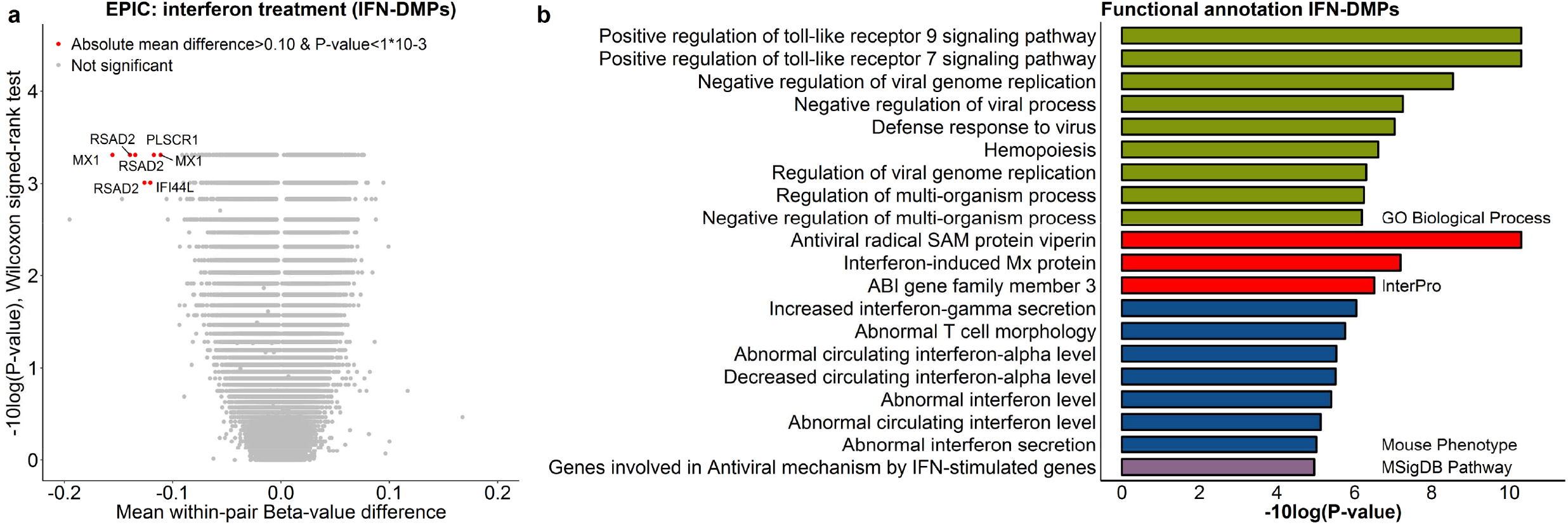
DNA methylation changes associated with interferon (IFN) treatment. (**a**) Results of the differential DNA methylation analysis including only the EPIC array data of the 12 pairs of which the MS-affected MZ co-twin was treated with IFN at the moment of blood collection. The volcano plot presents the P-values resulting from the non-parametric Wilcoxon signed-rank test versus the mean within-pair β-value difference for each CpG. Data were not adjusted for cell-type composition. Within-pair β-value difference (Δβ-value) = MS-affected IFN-treated MZ co-twin - clinically non-affected MZ co-twin. (**b**) Summary of the functional annotation analysis using GREAT^65^, on the 257 IFN-associated differentially methylated CpG positions (IFN-DMPs) (absolute mean within-pair β-value difference>0.05 and Wilcoxon signed-rank test P-value<0.001). Annotation terms are ranked according to their enrichment P-values: “GO Biological Process” terms (P-value<1*10^-6^) and the other presented terms (P-value<1*10^-5^).

### Glucocorticoid (GC) treatment induces hypomethylation at ***ZBTB16*** enhancer DMP

Among the MS-DMPs, the technically replicated *ZBTB16* DMP (cg25345365) had the largest effect size (~4%). This MS-DMP is located in an enhancer in intron 3 of *ZBTB16*, which encodes for a transcription factor also known as promyelocytic leukemia zinc finger (PLZF). ZBTB16/PLZF has been reported to be essential for NKT cell development^36^, and there is evidence that ZBTB16/PLZF contributes to T helper 17 (Th17) cell differentiation and phenotype maintenance^46^. However, *ZBTB16* has also been identified as a major glucocorticoid (GC) response gene being highly upregulated after GC exposure^47^, and several days of high-dose intravenous GC therapy is generally used to treat relapses in MS.

None of the MS-affected co-twins included in the array-based EWAS received GCs within three months prior blood collection, but 43 out of the 45 MS-affected co-twins have a GC treatment history, of which 14 received GCs within >3-12 months prior blood collection (minimal 3×1g and 6g on average). In those 14 pairs, the within-pair methylation differences at the *ZBTB16* DMP are significantly larger (more negative) compared to the pairs of which the MS-affected co-twin received GCs longer than 12 months ago (P=0.0004, **Fig. 4a** and **Supplementary Fig. 14**). This indicates that the strong association between the *ZBTB16* DMP and the MS phenotype is due to the GC treatment history in the MS-affected co-twins.

**Figure 4.**
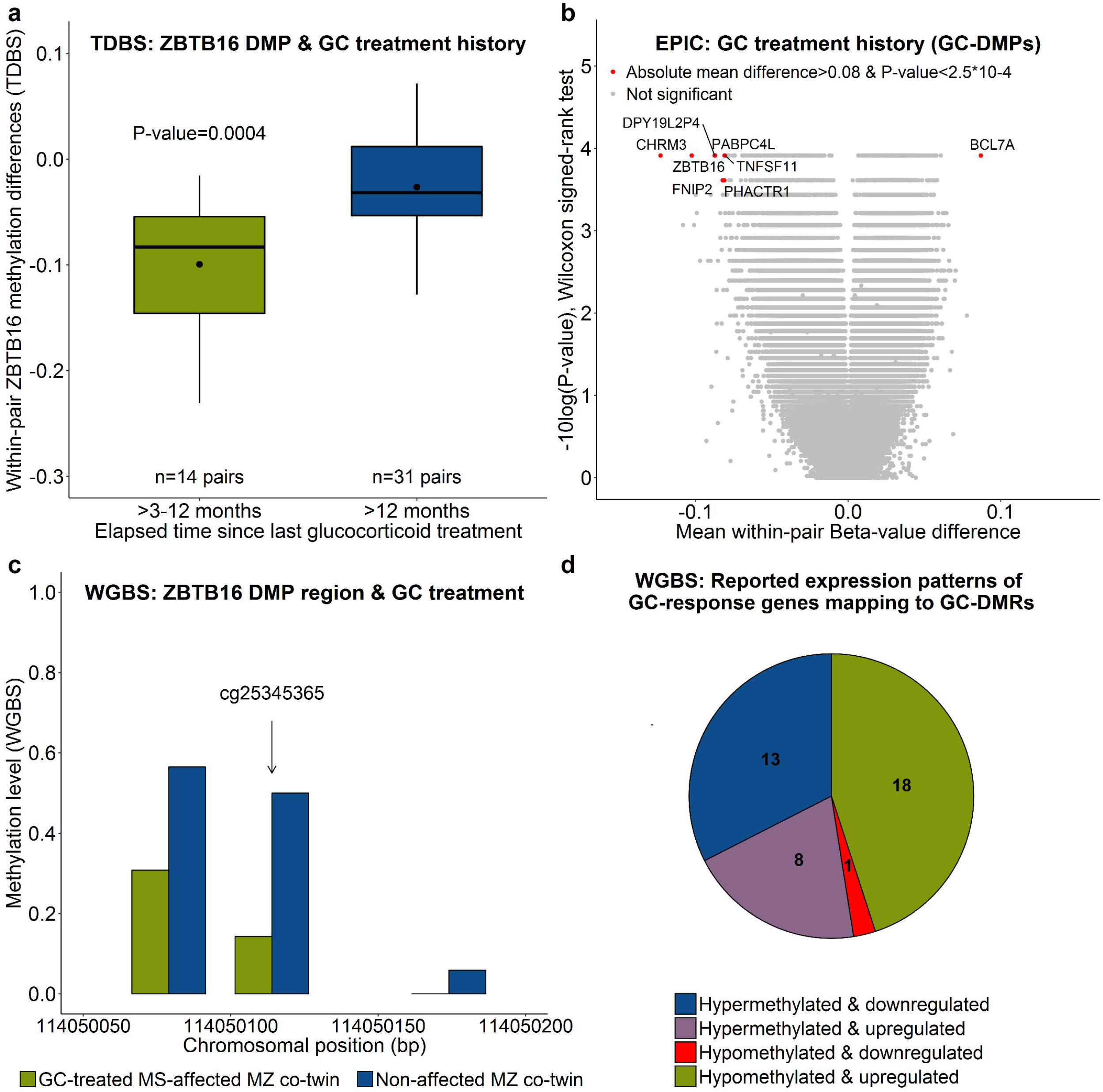
DNA methylation changes associated with glucocorticoid (GC) treatment (history). **(a)** Within-pair methylation differences of the *ZBTB16* DMP (cg25345365), determined using TDBS, in the 14 pairs of which the MS-affected co-twin received GCs within >3-12 months prior blood collection compared to the 31 pairs of which the MS-affected co-twin was treated with GCs more than 1 year ago. P-value = non-parametric Wilcoxon rank-sum test result. **(b)** Results of the differential DNA methylation analysis including only the EPIC array data of the 14 pairs of which the MS-affected co-twins received GCs >3-12 months prior blood collection. The volcano plot presents the P-values resulting from the non-parametric Wilcoxon signed-rank test versus the mean within-pair β-value difference for each CpG. Data were unadjusted for cell-type composition. Within-pair methylation/β-value difference = MS-affected MZ co-twin receiving GCs >3-12 months prior blood collection – clinically non-affected MZ co-twin. **(c)** Methylation level of the *ZBTB16* DMP (cg25345365) region determined using WGBS in CD4+ memory T cells of an MS discordant MZ twin pair of which the MS-affected MZ twin was treated with GCs at the moment of blood collection. Coverage at cg25345365 is >20 reads in each co-twin. **(d)** Methylation and reported expression patterns of the 41 GC-DMRs that overlap with GC-response (dexamethasone) genes recorded in the EMBL-EBI Expression Atlas (accessed May 2018). One GC-DMR was excluded, because it was reported to be down- and upregulated after dexamethasone treatment (**Supplementary Table 3**).

### GC-induced methylation changes are not widespread in GC-response genes

Then, the effect of recent GC treatment history on the PBMC methylomes was analysed by performing a pair-wise analysis including only the EPIC array data of 14 pairs of which the MS-affected co-twin received GCs within >3-12 months prior blood collection (**Fig. 4b**). In total, 320 potential GC-DMPs had an absolute mean Δß-value>0.05 and P<0.001, and were annotated to 279 genes. Of these, only five genes (1.8%), including *CCNA1, GMPR, ITGA6, LSP1* and *ZBTB16*, overlap with the 721 GC-response (dexamethasone) genes recorded in the EMBL-EBI Expression Atlas (accessed May 2018). Note that the other four MS-DMPs listed in **Table 1** were not among the 320 GC-DMPs.

To study acute effects of GC treatment, a WGBS analysis was performed on CD4+ memory T cells of an MS discordant MZ twin pair of which the MS-affected co-twin was treated with GCs at the moment of blood collection. As shown in **Fig. 4C**, the WGBS data confirmed the strong hypomethylation of the *ZBTB16* DMP (cg25345365) region in the GC-treated MS-affected MZ co-twin (36% methylation difference). In addition, 1424 other potential GC-DMRs were identified in the WGBS data, consisting of at least 2 CpGs, absolute mean methylation difference >0.25 and P<0.01. These GC-DMRs were annotated to 682 genes, and only 41 GC-DMRs overlap with 39 (5.7%) EMBL-EBI Expression Atlas reported GC-response genes (**Supplementary Table 3**), which represent potential GC-treatment epigenetic biomarkers. The majority of these 41 GC-DMRs were hypomethylated and the corresponding GC-response gene was recorded as upregulated due to GC treatment (**Fig. 4d**).

### *ZBTB16* enhancer DMP methylation correlates with global hypermethylation

Global DNA methylation levels of the PBMC-derived samples were assessed by TDBS of the repetitive elements *Alu*, human endogenous retrovirus type K *(HERVK)* and long interspersed nuclear element-1 *(LINE1).* Although *Alu* methylation correlated significantly with *LINE1* methylation (r=0.43, P=0.003), *Alu, HERVK* and *LINE1* methylation levels did not differ between the clinically non-affected and MS-affected co-twins (P>0.05). *Alu* and *HERVK* methylation were affected by cell-type composition differences, but the association also remained non-significant after adjusting for cell-type composition.

The volcano plots of the EPIC array data in **Supplementary Fig 1a** and **Fig. 2** are, however, slightly unbalanced, because 59.1% and 55.9% of the CpGs have a positive mean within-pair β-value difference in the analysis unadjusted and adjusted for cell-type composition, respectively. Since this is due to higher methylation values in the MS-affected co-twins, the EPIC array data does suggest a global hypermethylation in the MS affected co-twins. Although this global hypermethylation in the MS-affected co-twins was not significantly associated with GC treatment history (**Supplementary Fig. 15a**), the number of hypermethylated CpGs in the MS-affected cotwins was significantly correlated with the within-pair *ZBTB16* DMP methylation differences (r=-0.36, P=0.02) (**Supplementary Fig. 15b**).

### No evidence for within-pair copy number variations

Finally, discordant phenotypes within MZ twins can also be the result of within-pair copy number variations (WP-CNVs)^48^, therefore we checked the EPIC array data for CNVs. However, our analysis did not reveal evidence for chromosomal gains and losses within the MZ twin pairs. To illustrate, examples of copy number plots generated from the EPIC array data are presented in **Supplementary Fig. 16**.

## Discussion

Here we present the largest EWAS in MZ twins clinically discordant for MS to date. PBMC-based methylomes were generated using the EPIC array and revealed robust DNA methylation changes associated with IFN treatment, which demonstrates the sensitivity of the used technology. Overall, the PBMC-based methylomes of the 45 MS discordant MZ twins were highly similar, as the mean within-pair β-value differences were very small (<0.05). None of the MS-DMPs observed in our study have previously been reported by other EWAS for MS^19-25^. However, those MS EWAS studies observed much larger methylation differences and used selection criteria for MS-DMPs of absolute mean β-value differences >0.05^19^, or even >0.10^20-23^. Since these other studies used genetically unmatched cases and controls^19-25^, the large methylation differences observed by them might mainly be driven by genetic variation.

The most prominent MS-DMP in our EWAS was the technically replicated cg25345365 DMP in the transcription factor *ZBTB16*, which is also a GC-response gene that becomes highly upregulated after GC exposure^47^. Although nobody received GCs within three months prior blood collection, since 95% of the MS-affected co-twins have a GC treatment history and the healthy co-twins do not, GC treatment is a serious confounder. Our extensive analysis shows that the strong association between hypomethylation at the *ZBTB16* DMP and MS in our EWAS is due to the GC treatment history, which indicates that the generally used inclusion criterion of >3 months after completing GC treatment is not sufficient for epigenomic and transcriptomic studies in MS patients. **Supplementary Fig. 14** suggests a criterion of >1 year after GC treatment, but additional studies are required to address this issue properly. Our results might also have broader implications, because GCs are used in a wide variety of inflammatory and autoimmune diseases, but dosage and route of administration vary per disorder. The GC-glucocorticoid receptor (GCR) complex regulates transcription by binding to glucocorticoid response elements (GREs) in the genome^49^, and GC-GCR binding has been associated with DNA demethylation at enhancer elements^50^. This probably explains the hypomethylation observed at the *ZBTB16* DMP in our (GC-treated) MS-affected co-twins, since the DMP is located in an enhancer and is flanked (<100bp) by two consensus GRE downstream half-sites (TGTTCT) (**Supplementary Fig. 17**), which are believed to be sufficient for binding of the GC-GCR complex^49^. Although the precise demethylation mechanism remains unknown, data of Wiench et al.^50^ suggest that active demethylation takes place. Compared to the IFN analysis, we did not observe a very strong GC signature in the EPIC array data nor in the WGBS data, since in other common GC-regulated genes, such as *FKBP5, TSC22D3* and DUSP1^47,51,52^, no DNA methylation differences were observed. Altogether, our data reveals the *ZBTB16* DMP (cg25345365) as an epigenetic biomarker for GC treatment, and future studies have to assess its utility to predict clinical GC response in patients with inflammatory or autoimmune diseases receiving GC therapy.

Another prominent MS-DMP discovered by our EWAS is located in the promoter of *TMEM232* (cg27037608). Functional annotation analysis revealed that this region is enriched for MS-DMPs, suggesting that the *TMEM232* promoter harbours a DMR associated with the clinical manifestation of MS. Despite the small effect size (mean Δß-value=0.024), this DMR was technically replicated using TDBS, demonstrating that this represents a true effect. Furthermore, the *TMEM232* DMP was not among the GC or IFN-DMPs, indicating that the association with MS is probably not confounded by treatment history. Since the DMR is in close vicinity of the transcriptional start site, the methylation changes might be associated with expression changes, but this has to be evaluated. *TMEM232* is a member of the transmembrane (TMEM) protein family, that are predicted to be part of cellular membranes, including mitochondrial, endoplasmic reticulum, lysosomes and Golgi apparatus membranes^53^. Unfortunately, the function of *TMEM232* is to date unknown, and its possible role in the development of MS is unclear. However, variants in this gene have been associated with atopic dermatitis, allergic rhinitis and asthma in GWAS^30-32^, which might point towards a common immunologic pathway involving *TMEM232.* However, besides being biologically plausible, robust evidence that supports an association between atopic diseases and MS is thus far lacking^54,55^. At this point, further studies are needed to verify the association between the *TMEM232* DMR and MS.

We also carried out a WGBS analysis on CD4+ memory T cells of four MS discordant MZ twin pairs. This pilot did not reveal widespread global or site-specific methylation differences between the MS-affected and non-affected co-twins. Nevertheless, one potential MS-DMR was identified in an intronic regulatory region in the X-linked *FIRRE* gene (**Supplementary Fig. 13**), which encodes for a circular long non-coding RNA that has been reported to be involved in positioning the inactive X-chromosome to the nucleolus and to maintain histone H3K27me3 methylation^39,56^. Although our results are preliminary, MS is more common in women^3^ and a role of X-inactivation in the pathogenesis of MS has been proposed (reviewed by Brooks et al.^57^), therefore this DMR represents a possible candidate.

Our array-based EWAS allows us to detect small DNA methylation differences that are present in many patients, but MS is a heterogeneous disease and DNA methylation changes present in a few patients only might also contribute. In order to identify such rare methylation differences, we carried out a WP-DMR analysis using the EPIC array data. In total, 45 WP-DMRs were identified in 17 twin pairs, which indicates that WP-DMRs are quite common among MZ twins. Two WP-DMRs were present in two pairs, but not associated with the MS phenotype. The remaining 43 WP-DMRs were pair-specific, of which 27 showed an abnormal methylation level in the MS-affected co-twin. These WP-DMRs have not previously been associated with MS, although two WP-DMRs were related to genes encoding protocadherins that are involved in neuronal development^42^. Hence, a contribution of these WP-DMRs to the discordant phenotype cannot be excluded, but since they are pair-specific, these results should be interpreted very cautiously.

Due to the observed maternal parent-of-origin effect in MS and the reported involvement of several imprinted genes in the immune system, a role of genomic imprinting in the aetiology of MS has been suggested^15^. In this context, MZ twins are of particular interest, because MZ twins discordant for imprinting defects have relatively frequently been described^7,8^. Although we detected two WP-DMRs in imprinted regions *(SVOPL* and *HM13/MCTS2P)*, in both cases the aberrant methylation profile was observed in the non-affected co-twin. In addition, the EWAS did not reveal any MS-DMPs in imprinted genes. Consequently, our data does not support a contribution of genomic imprinting errors in the pathogenesis of MS.

Neven et al.^58^ reported hypermethylation of the repetitive elements *Alu, LINE1* and *Sat-α* in blood of MS patients compared to controls. Also Dunaeva et al.^59^ observed hypermethylation at a subset of *LINE1* CpGs in serum cell-free DNA of RRMS patients. Hence, we also assessed global methylation levels by TDBS of the repetitive elements *Alu, HERVK* and *LINE1*, but did not observe any significant differences between the MS-affected and non-affected co-twins. However, the unbalanced volcano plots in **Fig. 2a** and **Supplementary Fig. 1a** do suggest a slight global hypermethylation in the MS-affected co-twins. Also Bos et al.^19^ observed in their 450K data strong evidence for global hypermethylation in CD8+ T cells of MS patients, but not for CD4+ T cells or whole blood. Although in our data GC treatment history was not directly associated with global hypermethylation, we observed a significant association between increased within-pair *ZBTB16* methylation differences and the number of hypermethylated CpGs in the MS-affected co-twins. This might indirectly indicate that GCs also affect global DNA methylation levels, which might also explain the strong repetitive element hypermethylation in MS patients reported by Neven et al.^58^, who applied an inclusion criterion of only >1 month after GC treatment. Accordingly, additional studies are warranted to evaluate whether global hypermethylation in MS patients, observed by us and others^19,58,59^, is due to GC treatment history.

Although our discordant MZ twin design perfectly controls for genetic variation, our study also has limitations. MS discordant MZ twins are scarce and therefore it is not possible to control for treatment effects without losing statistical power. In addition, our data shows that besides shortterm, also medium-term treatment effects are measureable in immune cells, which makes recruitment even more challenging. Furthermore, some of the pairs included in this EWAS might get clinically concordant for MS in the future. However, at the moment the samples were taken the twins were clinically discordant for MS, enabling us to study DNA methylation differences that contribute to the discordant clinical manifestation of MS in MZ twins. For the EPIC array analysis we had only DNA extracted from PBMCs available, although it might be more informative to profile distinctive subtypes, such as CD4+ T cells, CD8+ T cells and B cells that are believed to be involved in the pathophysiology of MS^38,60^. However, a recent EWAS of 50 MZ twins discordant for type 1 diabetes that profiled CD4+ T cells, B cells and monocytes using the 450K array, observed only in the T cells one genome-wide significant DMP (mean Δβ-value=0.023)^61^. This might indicate that for detecting robust MS-DMPs it is required to immediately profile rare subpopulations such as Th1, Th17 and regulatory T cells, or to asses immune cells in the cerebrospinal fluid.

In conclusion, our EWAS shows that DNA methylation profiles in PBMCs of MZ twins clinically discordant for MS are overall very similar, and no evidence was found that genomic imprinting errors or CNVs explain discordance of MS in MZ twins. However, a DMR in the *TMEM232* promoter was identified as a candidate loci associated with the clinical manifestation of MS. In addition, epigenetic biomarkers for MS treatments were identified, revealing that besides shortterm also medium-term treatment effects are detectable in blood cells, which should be considered in epigenomic and transcriptomic studies. Altogether, we believe that this study represents an important first step in elucidating epigenetic mechanisms underlying the pathogenesis of MS.

## Methods

### Participants

Twins were recruited by launching a nationally televised appeal and internet notification in Germany with support from the German Multiple Sclerosis Society (DMSG, regional and national division). Inclusion criteria for study participation were met for MZ twins with an MS diagnosis according to the revised McDonald criteria or clinically isolated syndrome (CIS) in one co-twin only^62^. In total, 55 MZ twin pairs visited the outpatient department at the Institute of Clinical Neuroimmunology in Munich for a detailed interview and neurological examination. To confirm MS diagnosis, medical records including MRI scans from the patients’ treating neurologists were obtained and reviewed. For inclusion in the present analysis, peripheral blood mononuclear cells (PBMCs) had to be available of both co-twins, resulting in 46 MZ twin pairs. The pair that carries the Leber’s hereditary optic neuropathy-specific mutation m.11778G>A was not included in this analysis^6^. At blood collection, 23 MS-affected co-twins were treated with disease modifying treatments, including interferon-beta (IFN, n=12), natalizumab (n=5), glatiramer acetate (n=3), teriflunomide (n=1) and dimethyl fumarate (n=2). None of the MS-affected co-twins included in the array-based EWAS received glucocorticoids (GCs) within three months prior blood collection. The study was approved by the local ethics committee of the Ludwig Maximilians University of Munich and all participants gave written informed consent, according to the principles of the Declaration of Helsinki.

### DNA extraction and zygosity determination

PBMCs were isolated from whole blood using Ficoll density gradient centrifugation and DNA was extracted using the QIAamp DNA Blood Midi Kit (Qiagen, Hilden, Germany). Extracted DNA was treated with RNase A/T1 Mix (Thermo Scientific, Oberhausen, Germany), and subsequently purified using the Genomic DNA Clean & Concentrator™-10 Kit (Zymo Research, CA, USA). As previously described^6^, zygosity was confirmed by genotyping 17 highly polymorphic microsatellite markers and by next generation sequencing of 33 SNPs.

### Bisulfite treatment

From each sample, 500 ng DNA was bisulfite treated using the EZ DNA Methylation kit (D5002, Zymo Research), according to the manufacturer’s recommendations for the Illumina Infinium assay. The conversion reaction was incubated at 16 cycles of 95°C for 1 min and 50°C for 60 min. For all six MZ twin pairs that were processed in the first batch, the bisulfite controls present on the Illumina Infinium MethylationEPIC BeadChip (‘EPIC array’) showed suboptimal results, i.e. bisulfite conversion controls (I and II) showed moderate intensities at probes which should be at background level. However, the within-pair array-wide Pearson correlation coefficients for these MZ twin pairs were very high and ranged from 0.996 to 0.997, indicating high quality methylation data. We verified the conversion rate of these samples by targeted deep bisulfite sequencing (TDBS) of a 343-bp region amplified using non-bisulfite dependent primers (Chr1:202150908-202151251, forward 5’-TGGGGTAATGATGAGAGATGG-3’, reverse 5’-CTCTCTTTATTTCAAAACCCCCTA-3’). TDBS of these samples showed an average conversion rate of 98.6% (SD=0.37%) (minimal coverage >2000 reads), indicating that the EPIC array bisulfite controls are extremely sensitive. Hence, this EPIC array data was used in the downstream analysis. Nevertheless, the bisulfite treatment procedure was adapted by incubating samples in a programmable ThermoQ Metal Bath with heated lid (Bioer, Hangzhou, China) instead of a Eppendorf Mastercycler (Eppendorf AG, Hamburg, Germany), and TDBS in 16 samples revealed an average conversion rate of 99.7% (SD=0.10%). Accordingly, the other 40 MZ twin pairs were processed using the adapted bisulfite treatment and the EPIC array bisulfite controls showed normal intensities for those samples. Both members of a twin pair were always processed in the same batch.

### Infinium MethylationEPIC BeadChip assay

Genome-wide DNA methylation profiles of 46 MZ twin pairs clinically discordant for MS were generated using Illumina’s Infinium MethylationEPIC BeadChip assay (EPIC array) (Illumina, San Diego, CA, USA) at the Department of Psychiatry and Psychotherapy of the Saarland University Hospital. The assay allows determination of DNA methylation levels at >850,000 CpG sites and provides coverage of CpG islands, RefSeq genes, ENCODE open chromatin, ENCODE transcription factor binding sites and FANTOM5 enhancers. The assay was performed according to the manufacturer’s instructions and scanned on an Illumina HiScan. To avoid batch effects, both members of a twin pair were always assayed on the same array.

### Infinium MethylationEPIC BeadChip data processing and DMP identification

Raw EPIC array data were preprocessed using the RnBeads R/Bioconductor package^63^. Low-quality samples and probes were removed using the Greedycut algorithm, based on a detection P-value threshold of 0.05, as implemented in the RnBeads package. In addition, probes with less than three beads and probes with a missing value in at least 5% of the samples were removed. For each CpG site a β-value was calculated, which represents the fraction of methylated cytosines at that particular CpG site (0 = unmethylated, 1 = fully methylated). Subsequently, β-values were normalized using Illumina’s default normalization method. In total, methylation data of 849,832 sites (866,895 in total) were available for 45 MS discordant MZ twins. The relatively large number of excluded probes is due to inclusion of early access EPIC arrays, which have 11,652 probes less than the final release EPIC arrays. The EPIC array includes 59 SNP sites, which were used for quality control. All MZ twin pairs, except one, shared the same genotypes. The exceptional pair showed only a discordant genotype for SNP rs6471533. However, validation using targeted deep sequencing (TDS) revealed that both co twins have the same genotype for the rs6471533 SNP (**Supplementary Fig. 18**), which indicates a technical artefact in the corresponding EPIC probe rather than a true genetic difference.

To identify differentially methylated CpG positions (DMPs) a non-parametric Wilcoxon signed-rank test was carried out. For the MS EWAS, a significance level α<5*10^-6^ was considered suggestive and genome-wide significance was defined as false discovery rate (FDR) <0.05. All statistical analyses were performed in R^64^. A functional annotation analysis was performed using the Genomic Regions Enrichment of Annotations Tool (GREAT v3.0.0) with default settings and the EPIC array CpGs, that passed quality control, as background^65^.

### Power calculation

Since the power function of the Wilcoxon signed-rank test is difficult to express^66^, we used its closest parametric equivalent (paired t-test) to estimate the power of our MS EWAS. With a sample size of 45 twin pairs, >98% power is achieved to detect a mean β-value difference of at least 0.05 with a (genome-wide) significance threshold of 1×10^-7^, using a two-sided paired t-test and assuming a standard deviation of 0.0266 (which is the true standard deviation observed in our data). Details of this power calculation and calculations using smaller mean β-value differences are presented in **Supplementary Table 4**. The power analysis was performed using SAS University Edition.

### Estimation of cell-type composition

Cell-type composition of each PBMC sample was estimated with the reference-based method first published by Houseman et al.^67^, which uses DNA methylation reference profiles of individual cell-types to estimate the cell-type composition of each sample. Several reference-based deconvolution algorithms were compared, including the implementation of the Houseman algorithm in the *<u>minfi</u>* R/Bioconductor package^68^, and the standard constrained projection as well as the two non-constrained reference-based cell-type deconvolution approaches recently implemented in the EpiDISH R/Bioconductor package^69^. For a subset of samples (n=61) cell-type proportions determined using immunophenotyping were available, which showed the best correlation with the estimates provided by the *minfi* package. Accordingly, the *minfi* estimates were used to adjust the β-values for cellular composition using linear regression and the residuals were used for downstream analysis. To obtain interpretable adjusted β-values, the unadjusted mean β-value of each CpG site was added to the residuals. To check the quality of the adjustment, the adjusted β-values were used to recalculate the within-pair correlations. In the final regression model the proportions of the four major lymphocyte subtypes were included (i.e. CD4+ T, CD8+ T, CD19+ B and CD56+ NK cells). Myeloid cells (i.e. monocytes, neutrophils) were not included in the model, since immunophenotyping data showed that monocyte proportions were not properly estimated and inclusion resulted in severe adjustment bias in some samples. As a result, **Supplementary Fig. 19** shows that the overall within-pair correlations are, as expected, higher after adjusting for cell-type composition.

### Identification of within-pair differentially methylated regions (WP-DMRs)

To identify WP-DMRs in the EPIC array data, for each twin pair the β-value differences (Δβ-values) (adjusted for cell-type composition) per CpG were calculated (the 257 IFN-associated CpGs were excluded). To avoid false positives caused by single probes, WP-DMRs were defined as ≥3 CpGs having each an absolute Δβ-value >0.2 with a maximum 1 kb distance between neighbouring CpGs. To exclude regions that are characterized by overall variable methylation levels, WP-DMRs were only considered when the β-value of the aberrant methylated co-twin was more than three standard deviations away from the mean.

### Copy-number variation (CNV) analysis

CNV analysis with the EPIC array data was performed using the *conumee* R/Bioconductor package with default settings^70^. Individual profiles and output were manually assessed. To define chromosomal gains and losses within the MZ twin pairs, an absolute segment mean threshold ≥ 0.3 was applied.

### Targeted deep bisulfite sequencing (TDBS)

TDBS was used to validate DMPs resulting from the EPIC array analysis and to determine methylation levels of the repetitive elements *HERVK, LINE1* and Alu. Amplicons were generated on bisulfite-treated DNA using region-specific primers with TruSeq adaptor sequences on their 5’-ends (Illumina). Reaction conditions and primer sequences are described in **Supplementary Table 5.** Purified PCR products were quantified, pooled, amplified using index primers (five cycles), and sequenced in a 300-bp paired-end MiSeq run (Illumina). After demultiplexing, adaptor trimming and clipping overlapping mates, the resulting FASTQ files were imported into BiQ Analyzer HiMod^71^, to filter out low quality reads and call the methylation levels. Final coverage was >1500 reads/base.

### Targeted deep sequencing (TDS)

The rs6471533 SNP was genotyped using TDS (see **Supplementary Table 5** for reaction conditions and primer sequences). The workflow is similar as described for TDBS, except that genomic DNA was used and that the resulting FASTQ files were aligned to the reference sequence using Bowtie 2^72^. Subsequently, variants were called with SAMtools mpileup and variant information was extracted using filter pileup. Final coverage was >1500 reads/base.

### Whole-genome bisulfite sequencing (WGBS) and data pre-processing

WGBS was used to profile CD4+ central and effector memory T cells of four MS discordant female MZ twin pairs (mean age 43.3 years, discordant for MS >12 years, **Supplementary Table 6**). Of one pair the MS-affected co-twin was treated with GCs at the moment of blood collection (but never received any immune-modulating therapy), while the MS-affected co-twins of the other three pairs did not receive GCs or other immune-modulating therapies within at least 12 months prior blood collection.

Cryopreserved PBMCs were thawed and gently suspended in 10 ml of pre-cooled FACS buffer (PBS, 2% FCS), and centrifuged at 300 g for 10 min at 4°C. Then one additional washing step was performed. Cells were stained with the following monoclonal antibodies: CD3-AF700 (OKT3, eBioscience, Frankfurt, Germany); CD4-Pacific-Blue (S3.5, Molecular Probes, Invitrogen, Karlsruhe, Germany); CD8-PerCP (SK1, BioLegend, Fell, Germany); CD45RO-FITC (UCHL1, eBioscience) and CCR7-APC (3D12, eBioscience) on ice for 30 minutes. Cells were then sorted using a FACSAria Fusion flow cytometer (BD Biosciences, Heidelberg, Germany) to selectively collect antigen-experienced CD4+ T cells by excluding dead cells, naive CD45RA+CCR7+ T cells and CD8+ T cells.

WGBS libraries were prepared using a tagmentation-based protocol similar to that described by Weichenhan et al.^73^. Briefly, fresh frozen primary CD4 cell pellets (each 20,000-200,000 cells) were thawed in 50-100 μl of 1.1x TD buffer (Illumina) supplemented with 6 μl Protease (1 mg/ml; Qiagen) and incubated in a thermomixer at 55°C for 3 h followed by 20 min at 75°C. DNA was quantified using the Qubit HS-DNA kit (Thermo Fisher Scientific, Waltham, USA). From each sample the volume corresponding to 50 ng DNA was transferred in a new 1.5 ml tube and 1x TD buffer was added to a total volume of 47.5 μl. Then the DNA was tagged with 2.5 μl of Tn5 from the Nextera library preparation kit (Illumina) by incubation for 5 min at 55°C. After purification with the MinElute kit (Qiagen) and final elution with 10 μl EB buffer, gaps were repaired by adding 2 μl of 10x CutSmart buffer (NEB, Ipswich, USA), 3 μl of dNTPs (2.5 mM each), 5 U Klenow exo-(NEB) and incubation for 1 h at 30°C. Bisulfite conversion was performed with the EZ Methylation Gold Kit (Zymo Research) with a final 10 μl elution volume. Indexing library enrichment PCR was performed in 40 μl reactions with 1x HotStartBuffer (Qiagen), 0.25 mM of each dNTP, 0.3 μl ssDNA Binding Protein (Affymetrix, Santa Clara, USA), 100 nM of each primer (reverse primer contains sample-specific DNA barcode), 4 U HotStartTaq DNA polymerase (Qiagen) and 10 μl bisulfite-converted DNA. DNA was denatured at 95°C for 15 min, followed by 12 cycles of 30 sec at 95°C, 2 min at 53°C and 1 min at 72°C, and a final extension step of 7 min at 72°C. Reactions were purified using 0.8x volume AMPure XP Beads (Beckman Coulter, Brea, USA) and eluted in 10 μl Elution Buffer (Qiagen). Library fragment distributions were checked on the Agilent Bioanalyzer (Agilent, Santa Clara, USA).

The WGBS libraries were sequenced in a 100-bp paired-end HiSeq2500 run (Illumina) using custom sequencing primers. After adapter trimming using Trimmomatic v0.36^74^, the read pairs were aligned to the human reference genome (GRCh37) using *bwa-meth* v0.2.0^75^, which is a wrapper of the BWA-MEM1 alignment algorithm suited for bisulfite sequencing data. PCR duplicates were removed using the MarkDuplicates tool of the Picard suite v2.5.0-1 (http://broadinstitute.github.io/picard). Methylation levels of the CpG cytosines were determined using MethylDackel v0.2.1 (https://github.com/dpryan79/MethylDackel.git). Of both read mates 10 base pairs were disregarded from both read ends to eliminate the gap repair bias and methylation bias artefacts. The obtained BED files were loaded in the RnBeads package, which aggregated for each CpG the methylation information of both strands. The coverage statistics of the samples are summarized in **Supplementary Table 6**.

### DMR identification in WGBS data

To identify MS-associated DMRs (MS-DMRs), the WGBS data of all four pairs were analysed using the RnBeads package, in which for every CpG a paired t-test was performed. Only CpGs with a coverage ≥10 reads in all samples were included, resulting in methylation information of about 2.7 million CpGs (**Supplementary Table 6**). MS-DMRs were defined as ≥3 CpGs, each having P<0.05 and absolute mean methylation difference >0.2, and a maximum of 500 bp distance between neighbouring CpGs.

To identify glucocorticoid treatment-associated DMRs (GC-DMRs), the WGBS data of the pair with the GC-treated MS affected co-twin were analysed using DSS-single^76^, which is designed for detecting DMRs from WGBS data without replicates. To increase the quality of this single-replicate DMR analysis, only CpGs with a coverage ≥15 reads in both samples were included and the sex chromosomes were excluded, resulting in methylation data of up to 2.8 million CpGs (**Supplementary Table 6**). It has been reported that binding of the GC-glucocorticoid receptor (GCR) complex is rare within CpG islands and predominantly occurs at distal regulatory elements^50^. To detect DMRs in such CpG poor regions, the DSS-settings included a smoothing span of 100 bp, minimum DMR length of 25 bp with ≥2 CpGs and P<0.01. The absolute mean methylation difference had to be larger than 0.25, and to limit the number of false positives only GC-DMRs located in reported GC-response genes were considered. The GC and MS-DMRs were annotated using the ChIPseeker R/Bioconductor package (v1.14.2)^77^.

### Partially methylated domain (PMD) analysis in WGBS data

The WGBS data was segmented into PMDs, low methylated regions (LMRs) and unmethylated regions (UMRs) using the MethylSeekR R/Bioconductor package^78^. After filtering gaps annotated by UCSC^79^, the rest of the genome was designated as fully methylated regions (FMRs). As input for MethylSeekR the aggregated strand information per CpG was used, and the MethylSeekR settings included coverage ≥5 reads per CpG, 50% methylation and an FDR ≤ 0.05 for calling hypomethylated regions, resulting in a cut-off of ≥4 CpGs per LMR. For each segment, the methylation levels between the non-affected and MS-affected co-twins were compared using a paired t-test on the median weighted average methylation values. In addition, to assess the genome-wide PMD similarity across the eight samples, the genome was binned into 1kb windows, and each was annotated with 1 if the bin overlaps with a PMD and 0 otherwise. Based on this binarized matrix a hierarchical clustering was performed in R using “ward.D2” as agglomeration method and “euclidean” as a distance measurement. The very same procedure was performed for FMRs.

## Data availability

The epigenomic data has been deposited at the European Genome-phenome Archive (EGA, http://www.ebi.ac.uk/ega/), which is hosted at the EBI, under accession number EGAS00001003076 (submission in progress).

## Acknowledgements

We are grateful to all the twins, who participated in this study. We thank Jasmin Gries for performing MiSeq sequencing, Karl Nordström for pre-processing the reads, Kathrin Kattler for assisting tWGBS protocol optimization, and Katja Anslinger for zygosity determination. We acknowledge the use of the CF FlowCyt at the Biomedical Center of the Ludwig-Maximilians-Universität München. This work was supported by the Gemeinnützige Hertie Stiftung; German MS Foundation (regional and national division); German Research Council [SFB-TR 128 SyNergy]; Krankheitsbezogenes Kompetenznetz Multiple Sklerose, Cyliax Stiftung and the Verein zur Therapieforschung für MS Kranke.

## Author contributions

Study design: NS, LG, TK, RH, JW. Patient recruitment and care: LG, TK. Clinical data collection: LG. Experimental work and data collection: NS, GG, EB. Data processing: NS, PL. Data analysis: NS, AS. Supervising data processing and analysis: PL. Facilitating technical and material support: CP, DW, JW. Supervision: RH, JW. Manuscript writing: NS. Manuscript editing: all authors.

## Additional information

Supplementary information: This files contains 6 tables and 19 Figures

